# DIAmeter: Matching peptides to data-independent acquisition mass spectrometry data

**DOI:** 10.1101/2021.01.29.428872

**Authors:** Yang Young Lu, Jeff Bilmes, Ricard A Rodriguez-Mias, Judit Villén, William Stafford Noble

## Abstract

Tandem mass spectrometry data acquired using data independent acquisition (DIA) is challenging to interpret because the data exhibits complex structure along both the mass-to-charge (*m/z*) and time axes. The most common approach to analyzing this type of data makes use of a library of previously observed DIA data patterns (a “spectral library”), but this approach is expensive because the libraries do not typically generalize well across laboratories. Here we propose DIAmeter, a search engine that detects peptides in DIA data using only a peptide sequence database. Unlike other library-free DIA analysis methods, DIAmeter supports data generated using both wide and narrow isolation windows, can readily detect peptides containing post-translational modifications, can analyze data from a variety of instrument platforms, and is capable of detecting peptides even in the absence of detectable signal in the survey (MS1) scan.

## 1 Introduction

Data-independent acquisition (DIA) mass spectrometry (MS) is a powerful technique to study the proteome.^1,2^ Relative to data-dependent acquisition (DDA), DIA offers a broader dynamic range and more reproducible peptide detection. However, to achieve accurate quantification, DIA methods typically employ wide isolation windows to acquire a full mass range of fragmentation spectrum (MS2) data. As a result, multiple peptides are usually co-isolated and co-fragmented in the same MS2 scan. The resulting complex spectra cannot be effectively analyzed using conventional database search engines^3–7^ that follow a “one-peptide-per-spectrum” paradigm.

A common class of methods for analyzing DIA data relies on spectral libraries that store, for each charged peptide, fragmentation and retention time information.^8–13^ However, spectral libraries are expensive to produce, requiring considerable effort and resources devoted to sample preparation and data acquisition. Furthermore, the resulting libraries are typically not reusable across different laboratories or different instrument platforms.^14^ Some of these challenges can be addressed by generating spectral libraries *in silico*;^15–18^ however, the resulting predictions depend on the instrument and specific acquisition parameters and are typically less accurate than experiment-specific libraries.^19^ Furthermore, current state-of-the-art in *silico* prediction methods provide limited support for post-translational modifications (PTMs).^16^

An alternative class of library-free methods circumvent some of the challenges associated with generating and using spectral libraries.^12,20–22^ Library-free methods detect peptides from DIA data by using a peptide sequence database, rather than a library. The library-free approach is particularly valuable when production of a spectral library is impractical. These methods can be subdivided into *spectrum-centric* and *peptidecentric* approaches. Spectrum-centric methods, such as DIA-Umpire,^21^ extract coeluting and co-varying precursor and fragment ion groups into pseudospectra. These pseudospectra are then fed into a conventional database search engine designed for DDA-based peptide identification. The extraction of pseudospectra depends heavily on the quality of the precursor signals in the precursor (MS1) scans; hence, DIA-Umpire by design cannot detect peptides with undetectable precursor signals, which commonly arise due to limitations of intrascan dynamic range.^22^ Unlike spectrum-centric methods, peptide-centric methods such as PECAN^22^ query the DIA data for the best supporting evidence of detection for each peptide in the database. However, empirical evidence suggests that existing library-free methods are not as sensitive as library-based ones.^13,23^

Here we introduce DIAmeter, which detects peptides directly from DIA data without dependence on a spectral library (Figure 1). The input to DIAmeterincludes m/z-centroided DIA data and a proteome FASTA database. DIAmeter then searches the DIA data using a standard DDA search engine, allowing multiple peptide-spectrum matches (PSMs) per DIA spectrum. The PSMs are then augmented with auxiliary features describing various types of evidence supporting the detection of the associated peptide, including two different peptide-spectrum match scores, analysis of the precursor (MS1) signal, comparison of observed and predicted chromatographic retention time, and coelution of precursor and peptide fragment ions. PSMs are filtered using a weighted combination of these scores, and the PSM feature vectors are then processed by the Percolator machine learning post-processor^24^ to induce a ranking on peptides along with statistical confidence estimates, where highly ranked peptides are detected in the DIA data with stronger confidence.

**Figure 1:**
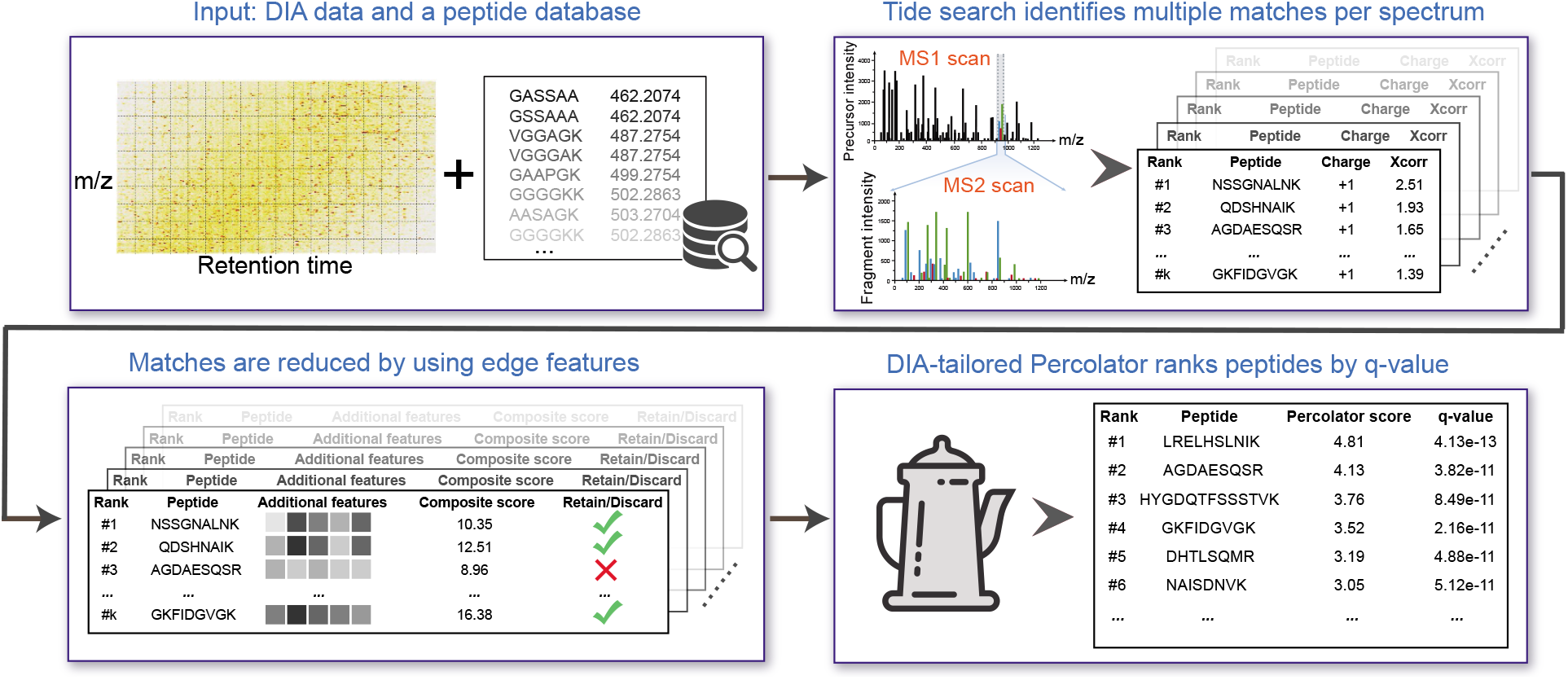
Overview of DIAmeter. The algorithm takes as input DIA data from a single run and a peptide sequence database. The output is a list of peptides, ranked by q-value, which is defined as the minimal FDR threshold at which that peptide is accepted. The Tide search step reports the top *k* peptide-spectrum matches (PSMs) per spectrum for charge states +1 through +5. In this work, *k* = 5. The edge features computed during composite scoring are described in Section 2.2.

Compared to existing DIA analysis methods, DIAmeteroffers four primary advantages (Table 1). First, DIAmeter by design can incorporate DIA data acquired using wide precursor isolation windows *(e.g.,* up to 25 Da). In this setting, the MS2 spectra are more complex because multiple peptides are more frequently co-fragmented. Second, DIAmeter can readily detect peptides with PTMs, as long as they are included in the database. Third, DIAmeter makes no assumptions about instrument-specific fragmentation patterns, instrument resolution, or patterns of background noise. Fourth, DIAmeter uses precursor information as a feature but does not require such evidence to detect a peptide. The open source, Apache licensed, Python version of DIAmeter is accessible at https://bitbucket.org/noblelab/diameter.

**Table 1:** Comparison of library-free peptide detection methods for DIA data.

|  | DIAmeter | Spectrum-centric<br>DIA-Umpire | Peptide-centric<br>PECAN | *in silico* library<br>Prosit+EncyclopeDIA |
| --- | --- | --- | --- | --- |
| Supports wide isolation windows | ✓ | ✓ |  |  |
| Detects PTMs | ✓ | ✓ | ✓ |  |
| Instrument independent | ✓ | ✓ |  |  |
| Detects peptides with undetectable precursor | ✓ |  | ✓ | ✓ |

## 2 Approach

The DIAmeter workflow (Figure 1) consists of the following steps.

1. Construct a bipartite graph between the observed DIA data and the list of theoretical precursors (i.e., charged peptides) in the database using a DDA search engine.
2. Compute a set of features for each edge in the graph, and eliminate some edges based upon a composite score aggregated from these features.
3. Run the remaining edges, with associated features, through a modified version of the Percolator algorithm.
4. Use target-decoy competition at the peptide level to estimate a q-value (i.e., the minimal FDR threshold at which a given peptide is accepted) for each ranked peptide.

These steps are described in more detail in Sections 2.1–2.4.

### 2.1 Bipartite graph construction

We conceptualize the DIAmeter approach using an undirected bipartite spectrum-to-precursor graph *G* = (*U,V,E*) that matches DIA spectra and a precursor database (Figure 1). Specifically, we define *U* to be the set of all MS2 spectra, each of which corresponds in a one-to-one fashion to a specific acquisition range in a single MS1 scan. For each spectrum *u ∈ U*, we let *t_u_* and *d_u_* denote its corresponding MS scan and acquisition range, respectively. The precursor database *V* is an unordered list of charged peptide sequences. For each precursor *v ∈ V*, we let *m_v_*, *c_v_* and *p_v_* denote its corresponding precursor m/z value, charge state, and peptide sequence, respectively. Conversely, *u ∈ U* and *v ∈ V* can also be indexed by *u_t_u_,d_u__* and *v_p_v_,c_v__*, respectively.

The edges *E* represent potential peptide-spectrum matches. Thus, for *v* ∈ *V* and *u* ∈ *U*, an undirected edge *e* = (*u, v*) ∈ *E* connects *u* and *v* if and only if the theoretical m/z value of the precursor v falls within a specified acquisition range *d_u_*. Because each spectrum *u ∈ U* may potentially be matched to a large number of precursors when the DIA isolation window is wide, we only keep the top *k* PSMs for each spectrum based upon their primary score. In this work, we use the SEQUEST XCorr score^25^ as the primary score, computed using the Tide search engine.^26^ We specify the parameter `--top-match 1000` and then keep the top-scoring *k* PSMs for each charge state *c* ∈ *C* where *C* = {1, 2,..., 5}. In this way, for each spectrum *u ∈ U*, at most |*C*| · *k* incident edges are added to the bipartite graph.

### 2.2 Calculation of edge features

For each edge *e* = (*u, v*) ∈ *E*, DIAmeter computes a set of eight features. These include three features generated by the Tide search engine—charge, delta_cn, delta_lcn—plus five edge features, described in detail below, each of which quantifies one type of evidence for or against the match between the spectrum u and the precursor *v*.

#### XCorr with Tailor calibration

The XCorr score is calibrated using the non-parametric Tailor calibration method,^27^ by specifying the parameter `--use-tailor-calibration T`. In brief, for each edge *e* = (*u,v*) ∈ *E*, the Tailor method calibrates the score XCorr_*u,v*_ by dividing by the 99th quantile of the observed, spectrum-specific score distribution. Specifically, say that spectrum *u* is scored with respect to *N* candidate peptides, each with charge state *c_v_*, during the database search. The resulting edge scores are sorted in decreasing order 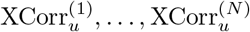, and the 99th quantile of this score distribution is obtained by identifying the edge score 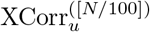 at position [*N*/100], where [·] denotes the round operator. The Tailor calibrated XCorr score is defined as

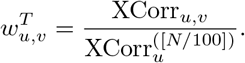

#### Precursor intensity rank

The observed precursor intensity can provide important evidence supporting the existence of a particular peptide in the sample, although as noted previously, lack of detectable signal in the precursor scan does not necessarily imply that the peptide has not been detected in the DIA data. For each edge *e* = (*u, v*) ∈ *E*, we denote 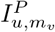 as the maximum observed precursor intensity at the precursor m/z value *m_v_* in the MS scan *t_u_*, within a user-specified tolerance. Using the raw precursor intensity directly is potentially problematic because the precursor intensity may exhibit biases specific to retention time, and because we don’t want a peptide with undetectable precursor signal to receive a score of zero. Therefore, we use the log-rank of precursor intensity instead of the raw intensity, represented as log 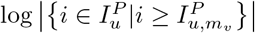, where 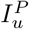 is the set of all observed peak intensities in the MS scan *t_u_*. In addition to the monoisotope of the precursor, we also consider *M* +1 and *M* + 2 isotopes. Therefore, the precursor intensity rank score is defined as

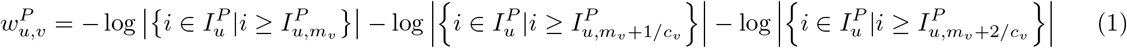

#### Fragment matching p-value

The fragment-matching p-value, analogous to the matching score in MS Amanda,^7^ measures how likely the theoretical fragments are to match the observed MS2 peaks by chance. Specifically, for each edge *e* = (*u, v*) ∈ *E*, by denoting *N_u,v_* and *N_v_* as the number of matched fragment peaks and all theoretical peaks, respectively, the fragment-matching p-value score is defined as the negative logarithm of the probability of matching at least *N_u,v_* peaks using the cumulative binomial distribution:

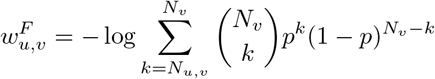

where *p_u_* is the probability to match one peak in u by chance. This value can be calculated as the fraction of the m/z range that is covered by the observed peaks. Analogous to the peak-picking in MS Amanda,^7^ the m/z range is divided into 10 equal length segments, and in each segment, the 10 most intense peaks are preserved. Let 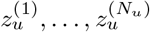 be the sorted m/z values, where *N_u_* = 100 is the total number of peaks remaining in the spectrum u. The fragment matching p-value *p_u_* is calculated as

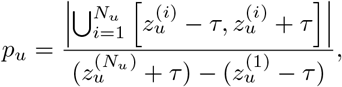

where τ specifies the matching tolerance.

#### Difference between observed and predicted retention time

An important factor to validate a candidate peptide is the difference between the observed and predicted retention times (RT). We use the scan number as a proxy for the observed RT, defining the set of observed RTs for all spectra as 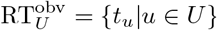. In parallel, we use the machine learning method DeepRT(+)o to define the set of predicted RTs for all precursors as 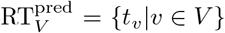, where *t_v_* indicates the predicted RT for the precursor *v ∈ V*. For each edge *e* = (*u, v*) ∈ *E*, the normalized observed and predicted RT can be represented as

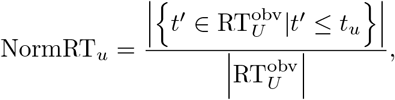

and similarly for NormRT_*v*_. Thus, the RT difference score is defined as

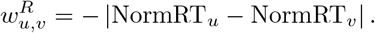

Note that DeepRT can only make RT predictions for unmodified peptides and peptides with methione oxidation or phosphorylation. For peptides with other types of modifications, DIAmeter uses the default value of 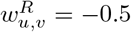.

#### Precursor and fragment coelution

Coelution of precursor ions and corresponding fragment ions is an important piece of evidence supporting detection of a specific peptide.^21^ In DIAmeter, this score is calculated as the normalized dot product between the corresponding elution profiles. Specifically, for each edge *e* = (*u, v*) ∈ *E*, the precursor elution profile spans nearby spectra within Δ*t* scan cycles (Δ*t* = 2 in our study), calculated as 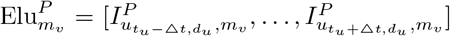, where 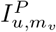 is the maximum observed precursor intensity at the precursor m/z value *m_v_* in the MS1 scan *t_u_* within some specified tolerance (see Section 3.4 for details). Similarly, by denoting 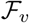 as the set of all fragment m/z values of the precursor *v*, for each fragment 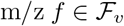, the fragment elution profile is calculated as 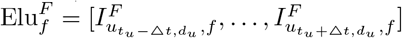, where 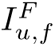 is denoted as the maximum observed fragment intensity at the fragment m/z value *f* in the MS2 scan *u*, also within some specified tolerance. We then define the set of normalized dot products between the precursor and its fragments as

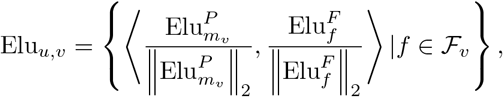

where 〈·,·〉 indicates the inner product operator. The final precursor/fragment coelution score 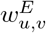 is the average of the top 3 values in Elu_*u, v*_.

All of DIAmeter’s hyperparameters are summarized in Supplementary Table S2.

#### 2.3 Edge re-scoring and filtering

Using the four scores described above, DIAmeter computes an aggregate score function and uses it to eliminate some edges in the bipartite graph. First, each score is linearly rescaled so that the 1st quantile and the 99th quantile map to the range [0,1]. Thereafter, the aggregated score is 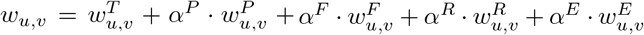, where the four non-negative hyperparameters *α^P^, α^F^, α^R^*, and *α^E^* balance the contributions from the corresponding score components. Subsequently, for each spectrum *u ∈ U* and charge state *c* ∈ {1,..., 5}, an edge is eliminated if its aggregated score is less than the aggregated score of the top-scoring edge according to XCorr. Formally, we denote 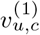 as the precursor corresponding to the maximum XCorr score in the set {*v* ∈ *V*|(*u, v*) ∈ *E,c_v_* = *c*}. The filtering step then discards an edge (*u, v*) if 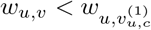.

### 2.4 Percolator tailored for DIA

Finally, all of the remaining edges are provided as input to the Percolator machine learning algorithm. Percolator’s semi-supervised algorithm learns to re-rank these edges by iteratively training a series of support vector machine classifiers, where negative examples are decoy edges and positive examples are high-scoring target edges.

Rather than using the FDR estimates provided by Percolator, DIAmeter calculates its own peptide-level FDR estimates. This choice is motivated by Percolator’s use of a spectrum-level target-decoy competition, in which only the highest-scoring PSM per spectrum is retained. In general, spectrum-level results are not useful for DIA analysis; therefore, we prefer to carry out the competition at the level of precursors, rather than PSMs. Hence, DIAmeter calls Percolator using the option `--tdc F`, which disables target-decoy competition. DIAmeter then selects the top-scoring edge per target-decoy pair; i.e., among all edges associated with a given precursor and that precursor’s corresponding decoy, only a single score is retained. The FDR estimate for a given score threshold *τ* is then calculated as usual:

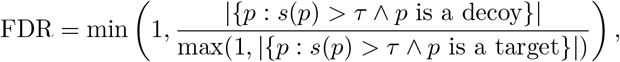

where *s*(*p*) is the Percolator score associated with precursor *p*.

One potential challenge associated with this approach to precursor-level FDR estimation is created by Percolator’s cross-validation procedure. This procedure equally divides the PSMs into three splits, from which the PSMs in two splits are used for training and the PSMs in the third split for prediction.^28^ This approach leads to a potential for “leakage” of information, in the case where, say, a decoy appears in the training set but its corresponding target is in the test set. To avoid this problem, we modified Percolator so that all PSMs related to the same target/decoy pair are assigned to the same cross-validation split.

## 3 Methods

### 3.1 Datasets

We used three different datasets to evaluate the performance of DIAmeter, in comparison to the three representative state-of-the-art methods mentioned in Table 1. The datasets were chosen to include different instrument platforms, wide and narrow isolation window sizes, fixed-window and variable-window acquisition schemes, and different dynamic ranges (Table S1). Full details about the three datasets are provided in Supplementary Section 3.2. For all three datasets, the database search was performed with the following parameters: fully tryptic digest with up to two missed cleavages, static modifications of carbamidomethyl for cysteines, and up to three variable modifications of oxidation of methionine.

### 3.2 Data

#### LFQBench

This is a widely used DIA benchmark for peptide identification and quantification,^29^ which consists of lysate mixtures from human, yeast, and *E. coli* mixed in predefined proportions. We carried out the evaluation by using LFQBench samples acquired on two mass spectrometers (TripleTOF 5600 and TripleTOF 6600) with two acquisition schemes (32 fixed windows of 25 m/z effective precursor isolation and 64 variable windows optimized for roughly equal precursor partition) over a 120-minute gradient. The data as .wiff files, as well as the FASTA database combining human, yeast, and *E. coli* proteins, were downloaded from the ftp://ftp.pride.ebi.ac.uk/pride/data/archive/2016/09/PXD002952.

#### Plasma

This dataset is derived from a large cohort designed to assess the heritability and environmental effects on blood plasma protein abundance.^30^ The data was acquired on a TripleTOF 5600 mass spectrometer with 32 fixed windows of 25 m/z effective precursor isolation over a 120-minute gradient. The data as .wiff files, as well as the human protein FASTA database, were downloaded from ftp://ftp.pride.ebi.ac.uk/pride/data/archive/2015/01/PXD001064.

#### OxMetYeast

This dataset consists of three mass spectrometry experiments where different isolation window sizes were explored in combination with gas phase fractionation, using a yeast lysate with selective oxidation of methionine. The resulting DIA data, as .raw files, include (1) one run covering the 400-1200 m/z mass range acquired using 20 m/z isolation windows; (2) two runs acquired using 10 m/z isolation windows to cover 400-800 and 800-1200 m/z mass ranges; (3) four runs acquired using 5 m/z isolation windows covering 400-600, 600-800, 800-1000, and 1000-1200 m/z ranges, respectively. The dataset has been deposited to the PRIDE repository^31^ under data set identifier XXX.^1^

To prepare the yeast sample, 25 *μ*g of tryptic yeast peptides, prepared as described by Studer *et al.*,^32^ were resuspended in 30 *μ*l of oxidizing solution containing 100 mM DMSO, 5% acetonitrile and 1 M HCl, and reacted at room temperature for 20 minutes. Peptides were then diluted two-fold with 0.1% TFA and desalted over a 2-layer C18 Stage tip (Empore, 3M),^33^ that had been previously conditioned with 20 *μ*l methanol, followed by 20 *μ*l 50% acetonitrile 0.5% acetic acid, and 20 *μ*l 0.1% TFA. Peptides on the Stage tip were washed 3 times with 20 *μ*l of 0.1% TFA and eluted with 50% acetonitrile 0.5% acetic acid. Eluted peptides were then lyophilized.

Lyophilized peptides were resuspended in 50 *μ*l of 5% ACN, 5% formic acid and subjected to liquid chromatography coupled to tandem MS. A total of 1.5 *μ*l of sample was loaded onto a 100 *μ*m × 35 cm fused silica capillary tubing packed with 1.9 *μ*m ReproSil-Pur C18 (Dr. Maisch) reversed-phase material and separated using a gradient of 5% to 36% acetonitrile in 0.125% formic acid delivered at 300 nl/min over 73 minutes on an EASY-nLC 1200 nano-flow UHPLC system (Thermo Scientific). Peptides were online analyzed on an Orbitrap Eclipse Tribrid MS (Thermo Scientific) with a total 90-minute acquisition time. All methods consisted of an MS1 scan at 60K resolution every 3 seconds with 100% normalized AGC target and a maximum injection time of 50 ms followed by DIA MS2 scans events acquired at 30K resolution with a maximum injection time of 54 ms and 4e5 target AGC.

#### File conversion

The .raw files generated by Orbitrap Eclipse Tribrid were converted to .mzXML (used in DIAmeter and DIA-Umpire) and .mzML (used in EncyclopeDIA) files by using the msconvert provided with ProteoWizard.^34^ The .wiff files generated by TripleTOF instruments were converted to .mzXML and .mzML files by using the qtofpeakpicker provided with ProteoWizard,^34^ with the default options: `--resolution 20000 --area 1 --threshold 10 --numberofpeaks 0`.

#### Noise reduction

DIAmeter applies a noise reduction strategy^35^ to TripleTOF 5600 data by following the assumption that peaks occurring in a single scan, without any corresponding peaks in preceding or succeeding scans, are noise. Specifically, the MS2 peak is eliminated if neither of the adjacent scans contains the same peak within a specified tolerance. For the Orbitrap and TripleTOF 6600 instruments, noise reduction is not applied due to their high signal-to-noise ratio.

### 3.3 Consistent FDR estimation across methods

One significant challenge when comparing different peptide detection methods is that each method chooses to estimate FDR in its own fashion. Therefore, an improvement in statistical power for method *A* over method *B* may arise either because A outperforms *B* or because *B* makes more conservative FDR estimates than *A*.

To address this challenge, we use a consistent “pseudo-target” scheme to evaluate all methods. The approach, similar to previously described “entrapment” strategies,^36^ is as follows. First, given a target peptide database, a decoy peptide database is generated of the same size. Both sets of sequences are provided to the peptide detection method, but both are labeled as targets. The method then uses its own internal procedure to generate a decoy database, search the concatenated target-decoy database, and induces a ranking on the peptide as output. We discard the reported FDR and instead use the pseudo-targets to estimate FDR, as in Section 2.4. The inclusion of pseudo-targets leads to a slight loss of statistical power but is only done for evaluation purposes.

### 3.4 Benchmark settings

#### DIAmeter

The first step of DIAmeter involves a database search that considers, for each spectrum, all possible precursors whose corresponding m/z value falls within the DIA isolation window. To do so, the precursor mass for each DIA spectrum is calculated based on the center m/z value of its corresponding isolation window, and searched using the Tide search engine^26^ with the parameter `--precursor-window-type mz --precursor-window <isolation window radius>`. Afterwards, the bipartite spectrum-to-precursor graph is constructed based on the top *k* = 5 PSMs for each spectrum and for each charge state. In the subsequent feature calculation step, a precursor and fragment mass tolerance of 10 ppm is used for all Orbitrap datasets, and 30 ppm for all TripleTOF datasets.

The DIAmeter hyperparameters are selected via cross-validation over the 8 experiments involved in the evaluation. For each experiment regarded as the test set, we treated the remaining 7 experiments as the training set and select the best-performing hyperparameters from a grid in which each of the *α* values (*α^P^, α^F^, α^R^, α^E^*) can take any value in the set {0, 0.05,0.1,0.2,0.4,0.8,1.6, 3.2, 6.4,12.8}. The performance of the chosen hyperparameters is evaluated by the number of accepted peptides at a 1% peptide-level FDR.

#### DIA-Umpire

For DIA-Umpire, the preprocessed .mzXML files were analyzed by the signal extraction module of DIA-Umpire (version 2.0) to generate pseudo pectra in MGF format, at all quality tiers, which were subsequently searched using the Tide search engine.^26^ We kept the search settings as close as possible to the DIAmeter settings to facilitate a fair comparison. The complete list of DIA-Umpire parameter settings is provided in Supplementary Table S3.

#### PECAN

We compared against Walnut, a re-implementation of PECAN provided by EncyclopeDIA (version 0.9.0),^13^ with similar setting as DIAmetersettings. Full parameters are in Section 3.2.

#### Prosit+EncyclopeDIA

For EncyclopeDIA,^13^ we used an *in silico* library generated by Prosit.^17^ The fragmentation prediction in Prosit is adjusted based on the optimal normalized collision energy setting to account for DIA-specific fragmentation, as suggested previously.^19^ For the Prosit-generated spectral library, by design peptides with length in the range of [7, 30] and charge state *C* = {2, 3} are considered.

## 4 Results

### 4.1 DIAmeter confidently detects more peptides

We systematically evaluated the performance of DIAmeter relative to DIA-Umpire, PECAN and Prosit+-EncyclopeDIA by counting the number of distinct peptides detected in various datasets at a 1% FDR threshold (Figure 2). For this analysis, FDR is estimated using the pseudo-targets described in Section 3.3.

**Figure 2:**
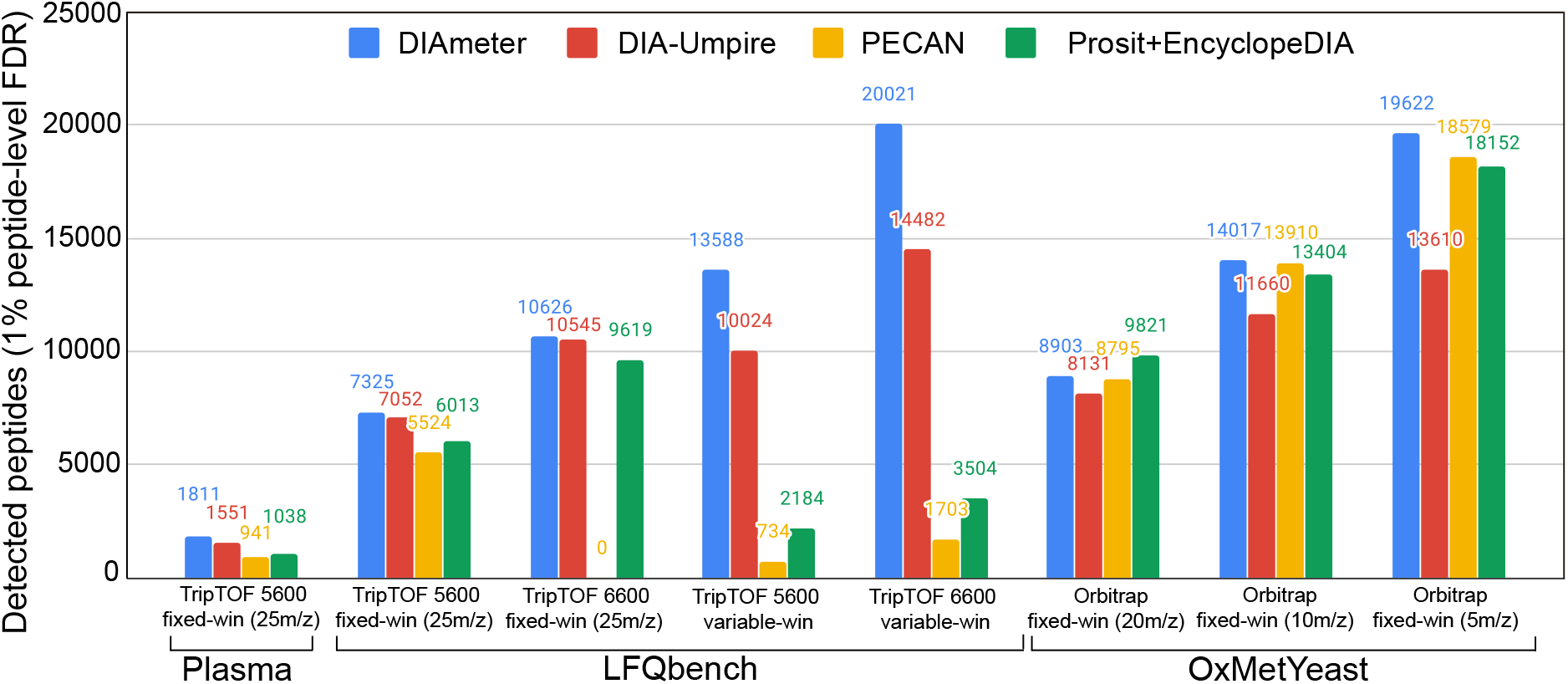
Performance comparison across various methods. The performance is evaluated by the number of detected peptide, at a 1% peptide-level FDR, in eight experiments from three different datasets.

We first focused on the LFQBench dataset, which allows us to consider how different instrument platforms impact the sensitivity of peptide detection. This dataset is comprised of four mass spectrometry experiments acquired on two different types of mass spectrometer with two different acquisition schemes. At every setting, DIAmeter consistently detects more peptides than the three competing methods, with an average increase of 14.6% relative to DIA-Umpire, 77.7% relative to PECAN, and 48.5% relative to Prosit+EncyclopeDIA. The relatively large performance improvement relative to PECAN and Prosit+Encyclopedia may arise because these methods are tailored for Orbitrap instruments. For example, the Prosit prediction model was trained exclusively on Orbitrap data.^17^

Not surprisingly, the TripleTOF 6600 provides improved peptide detection over its predecessor, the 5600, consistently across the four analysis methods. For example, for DIAmeter switching from the TripleTOF 5600 to the 6600 yields an average increase of 46.2% in the number of detected peptides. The only notable exception to this trend is PECAN, where no peptide is confidently detected within 1% FDR threshold on the TripleTOF 6600. We hypothesized that this failure might be due to the large size of the peptide database. This hypothesis is supported by the observation that, when we randomly discarded 70% of the targets and their corresponding decoys from the database, the number of detections increased from 0 to 844.

Next, we tested how the performance of each of the four methods changes as we vary the size of the precursor isolation window. In general, employing a wide isolation window drastically increases the complexity of the resulting spectra and in turn decreases detection sensitivity.^22^ To test how isolation window size impacts detection performance, we used the OxMetYeast dataset, which consists of three mass spectrometry experiments acquired using isolation windows of 20 m/z, 10 m/z, and 5 m/z. As expected, all four methods detect more peptides as the window size decreases. For example, DIAmeter detects 8903, 14017, and 19622 peptides, respectively. On average across the four methods, the number of detected peptides increases by 23.5% when we switch from 20 m/z to 10 m/z and by 32.6% when we switch from 10 m/z to 5 m/z. Among the four methods, DIAmeter performs best overall for narrower windows, and second best (behind Prosit+EncyclopeDIA) for 20 m/z windows.

Finally, to investigate the robustness of the four analysis methods to samples showing a large dynamic range of peptide abundance, we used a human plasma dataset acquired on a TripleTOF 5600. DIAmeter detected 1811 unique peptides in this dataset, outperforming DIA-Umpire, PECAN, and Prosit+-EncyclopeDIA, which reported 1551, 941, and 1038 unique peptides, respectively. Thus, overall, DIAmeter shows robust peptide detection performance across a variety of datasets, in comparison with three methodologically distinct methods for DIA analysis. On the other hand, investigation of detection rates at other FDR thresholds (Figure 3) suggests that, in some cases, the relative performance among methods depends on the selected FDR threshold. In particular, in most tripleTOF experiments, DIAmeter consistently performs well across a range of q-value thresholds, whereas in Orbitrap experiments, DIAmeter performs best for q-values ≤ 0.01.

**Figure 3:**
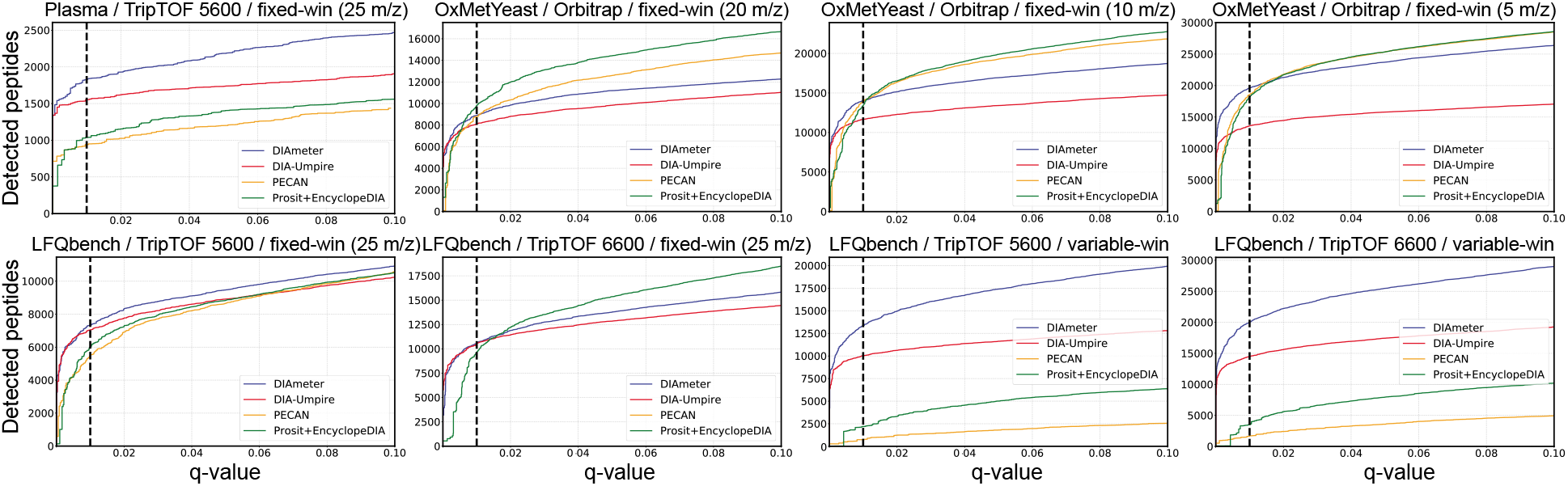
Performance comparison as a function of FDR threshold. Performance of each method is evaluated by the number of detected peptides as a function of peptide-level q-values in eight experiments from three different datasets.

### 4.2 DIAmeter can detect peptides with undetectable MS1 signal

A central challenge in the analysis of any protein tandem mass spectrometry data set is the huge dynamic range of protein abundances in many complex biological samples. In DIA analysis, this variability in protein abundance means that some peptides without detectable precursor signals may still be identifiable on the basis of their fragmentation scans.^22^ We hypothesized that these low abundance peptides may be particularly problematic for a method like DIA-Umpire, which is designed to construct pseudo-spectra on the basis of precursor signals.

To test this hypothesis, we compared the peptides detected by DIA-Umpire and DIAmeter in three representative datasets acquired on Orbitrap, TripleTOF 5600 and TripleTOF 6600 mass spectrometers. We reasoned that if a peptide exhibits low precursor signal, then DIA-Umpire is unlikely to construct a pseudo-spectrum that corresponds to that peptide. Therefore, for each dataset, we searched for peptides that are detected with high confidence (<1% FDR) by DIAmeterbut for which DIA-Umpire produces no corresponding pseudo-spectrum. In this analysis, we say a peptide corresponds to a pseudo-spectrum if the two m/z values are within the user-specified tolerance (in units of ppm) and if the retention time associated with the pseudo-spectrum is within ±15 s of the peak peptide detection. We further classified the detected precursors with no corresponding pseudo-spectra into two types: those for which no m/z match is available at any retention time (Type 1), and those for which a match exists, but only outside of the ±15 s RT window (Type 2).

Our analysis shows that DIAmeter detects many Type 1 and Type 2 precursors, regardless of the mass spectrometer type. In the data acquired on an Orbitrap (OxMetYeast with 20 m/z isolation window), among 11,355 precursors detected by DIAmeter with high confidence, 1064 (9.4%, 526 Type 1 and 538 Type 2) do not coincide with any pseudo-spectra extracted by DIA-Umpire at any quality tier. Among the 538 Type 2 precursors, 477 are identified by DIA-Umpire as different precursors with low confidence (average q-value 47.4%). Analogously, in the experiments acquired by the TripleTOF 5600 and TripleTOF 6600 (LFQBench with fixed window scheme), among 9968 and 14,476 confidently detected precursors reported by DIAmeter, 117 (1.2%, 29 Type 1 and 88 Type 2) out of 9968 and 205 (1.4%, 32 Type 1 and 173 Type 2) out of 14,476 do not coincide with any pseudo-spectra extracted by DIA-Umpire at any quality tier, respectively. Furthermore, 87 out of 88 and 164 out of 173 detected Type 2 precursors are identified by DIA-Umpire as different precursors with low confidence (average q-value 62.3% and 58.5%) in two settings, respectively.

Finally, we showcase confidently identified peptides detected in different instrument platforms (Figure 4). These peptide exhibited the lowest q-value among all peptides that lacked any corresponding pseudo-spectra. It is worth mentioning that DIAmeter can confidently detect precursors with both M+1 and M+2 isotope signals but without any monoisotopic signal (Figure 4(b)), which might be caused by glutamine deamidation. Thus, we demonstrated the credibility of peptides without detectable precursor signals, in terms of the observed fragment ions in the matched MS2 spectrum as well as the coeluting ion chromatograms.

**Figure 4:**
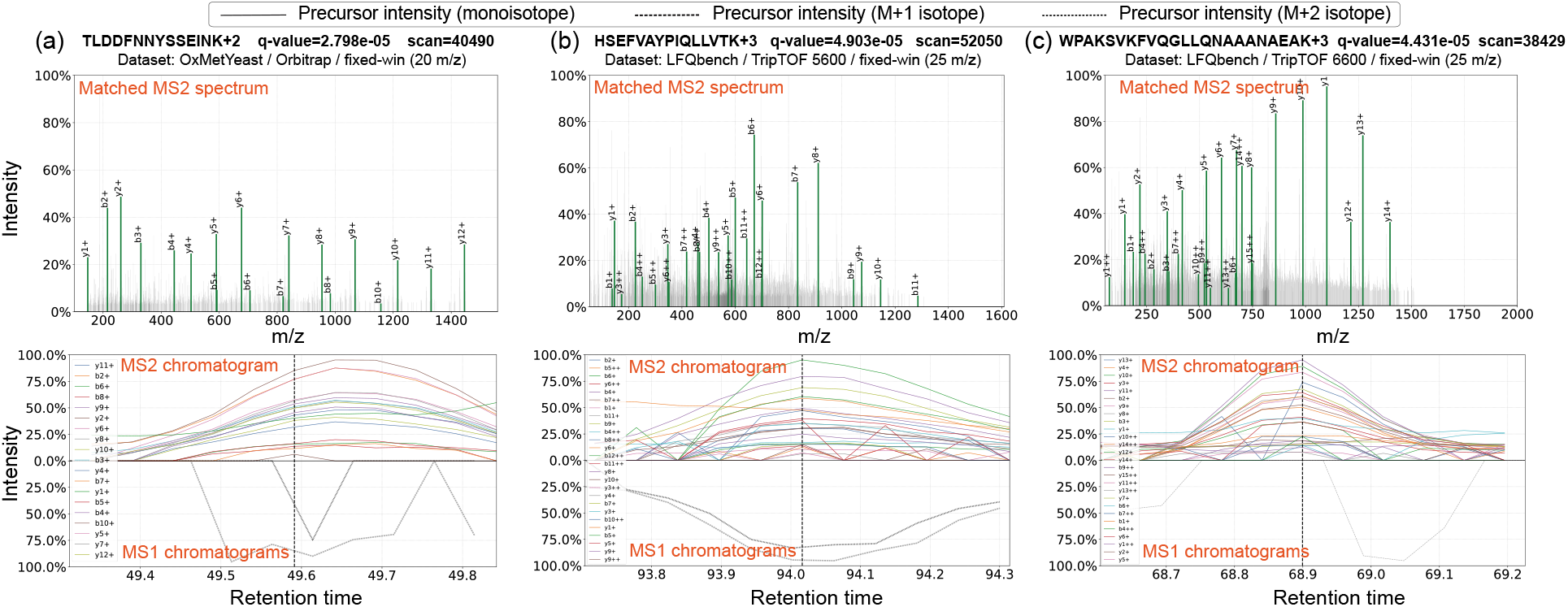
Examples of peptides confidently detected by DIAmeter. The figure shows peptides detected on different mass spectrometers, in terms of the observed fragment ions matched in the MS2 spectrum as well as the MS1 chromatograms in ±5 scans.

### 4.3 Necessity of each DIAmeter component

A key component of DIAmeter is its use of a variety of sources of evidence to support detection of a single peptide, including the quality of the match between the observed and theoretical spectra, properties of the precursor isotope distribution, the difference between observed and predicted retention time, and properties of the peptide chromatogram. To evaluate the extent to which each of these features contributes to the peptide detection, we ran an ablation study, in which we modified DIAmeter to include various subsets of the full collection of score features. In particular, we considered 10 variants of DIAmeter: five in which we eliminate a single feature (the XCorr score, the fragment matching p-value, the precursor intensity, the RT difference score, and the chromatographic score), and five in which we retain only a single feature. In each case, we run DIAmeter using the reduced features for edge re-scoring, edge filtering, and Percolator postprocessing. We carried out this analysis on the OxMetYeast (20 m/z window) and LFQBench experiments, representing three different instrument platforms.

The results of this analysis (Figure 5) suggest that all five features contribute positively to DIAmeter’s performance. In particular, the Tailor calibrated XCorr score appears to be the most informative feature, from both the inclusion and exclusion perspective, suggesting that peak matching provides the most important evidence in DIA data. Conversely, the fragment matching p-value appears to be the least informative feature on its own. Nonetheless, in combination with other features, inclusion of the fragment matching p-value facilitates better peptide detection, particularly on TripleTOF 6600 instruments.

**Figure 5:**
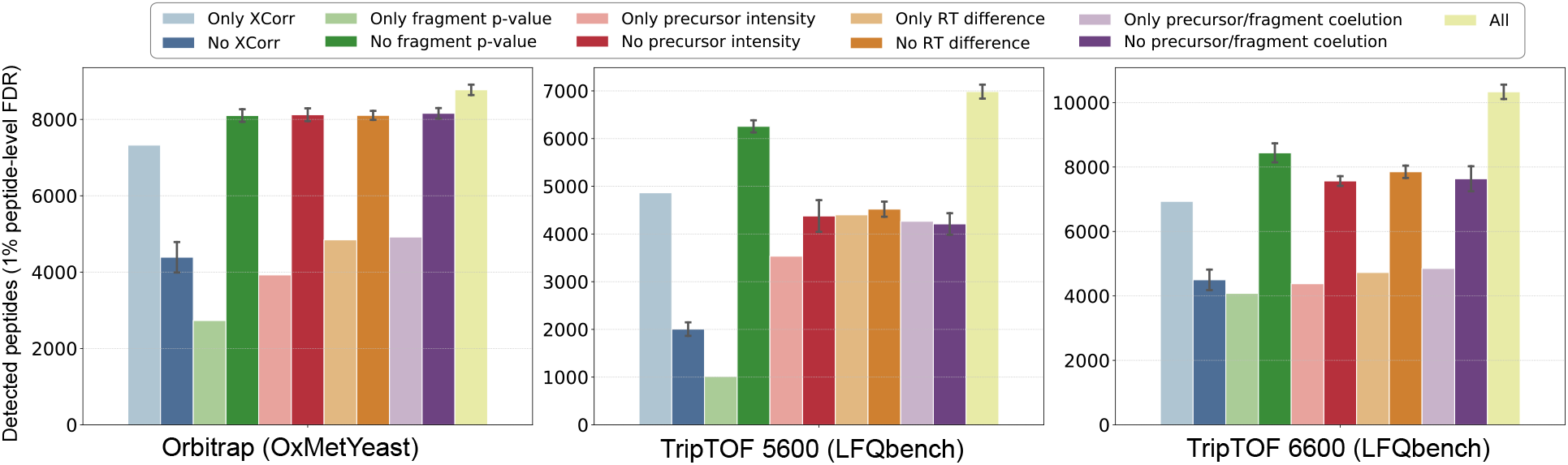
Evaluation of the marginal contribution of each edge feature on different instrument platforms. Each bar shows the number of peptide detected by a modified variant of DIAmeter. Modifications use either individual edge features or all but one of the edge features.

It is worth mentioning that the utility of some features appears to vary by instrument. For example, exclusion of the precursor/fragment coelution score or the RT difference score makes a negligible contribution to peptide detection for data generated on the Orbitrap, but removing this feature from DIAmeter compromises performance on the TripleTOF instruments. Finally, the inclusion of 5 features in DIAmeter does not fully capture the sample/instrument/acquisition variations. It would be an interesting pursuit in the future to extend DIAmeter to incorporate richer information such as sample complexity, dynamic range, mass accuracy, and m/z windows

### 4.4 Comparison of FDR control methods

We implemented the pseudo-targets FDR estimation strategy described in Section 3.3 in order to ensure a fair comparison of different DIA analysis methods; however, this approach also allows us to compare the different FDR estimation procedures employed by these methods. To do so, we plot the nominal q-values reported by each method against the q-values estimated using the hidden pseudo-targets. The results on three different instrument platforms (Figure 6) suggest that, in general, three of the DIA analysis methods tend to be conservative, in the sense that the nominal q-values are larger than the pseudo-target q-values. The exception is Prosit+EncyclopeDIA, which yields the reverse direction for two out of three datasets. Note that these results cannot conclusively demonstrate either the validity or invalidity of the nominal q-values because the FDR is defined as the *expectation* (roughly, the average over all possible datasets) of the proportion of false discoveries at a given score threshold. Nonetheless, the results are suggestive that some of these methods (in particular, DIA-Umpire and PECAN) could potentially gain statistical power simply by improving their methodology for FDR estimation.

**Figure 6:**
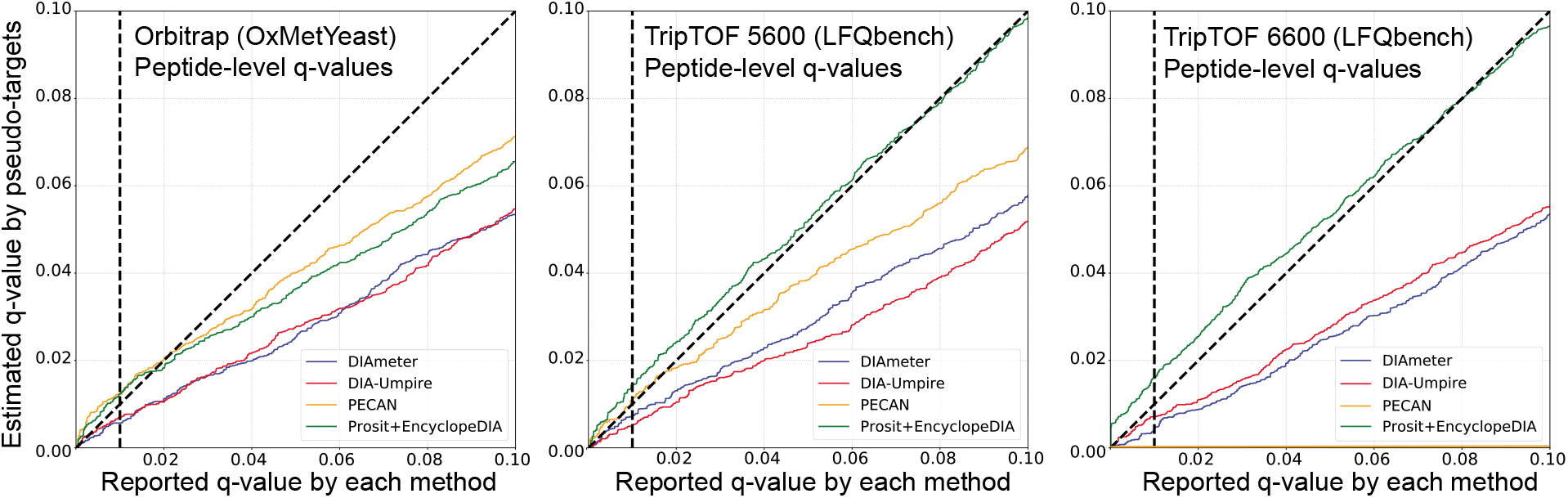
Comparison of FDR control methods. The figure plots the nominal peptide-level q-values reported by each method versus the corresponding q-values estimated by using pseudo-targets, on three datasets acquired by different instrument platforms.

## 5 Discussion

In this work, we introduced a novel and versatile method, DIAmeter, to detect peptides directly from DIA data without dependence on a spectral library. Compared to existing DIA analysis methods, DIAmeter offers several advantages. First, DIAmeter by design can incorporate DIA data acquired using wide precursor isolation windows. Second, DIAmeter is based upon a conventional DDA database search engine, which is readily able to detect peptides with PTMs. Third, DIAmeter works well on a variety of instrument platforms, because the method makes no assumptions about instrument-specific fragmentation patterns, instrument resolution, or patterns of background noise. Forth, DIAmeter by design can detect peptides with weak or even undetectable precursor signals.

This work points to several promising directions for future research. One possible extension is to accelerate DIAmeter by converting the matching paradigm from our current spectrum-to-peptide matching approach to the spectrum-to-peptides matching approach employed by MSFragger,^37^ with the aid of a fragment index. Another promising direction could be customizing the spectrum matching score function to incorporate prior knowledge about peptide fragmentation.^38^

## Acknowledgments

This work was supported by National Institutes of Health awards R01 GM121818, R35 GM119536, and P41 GM103533, as well as a grant from the Keck Foundation.

## A Supplement

**Table S1:**
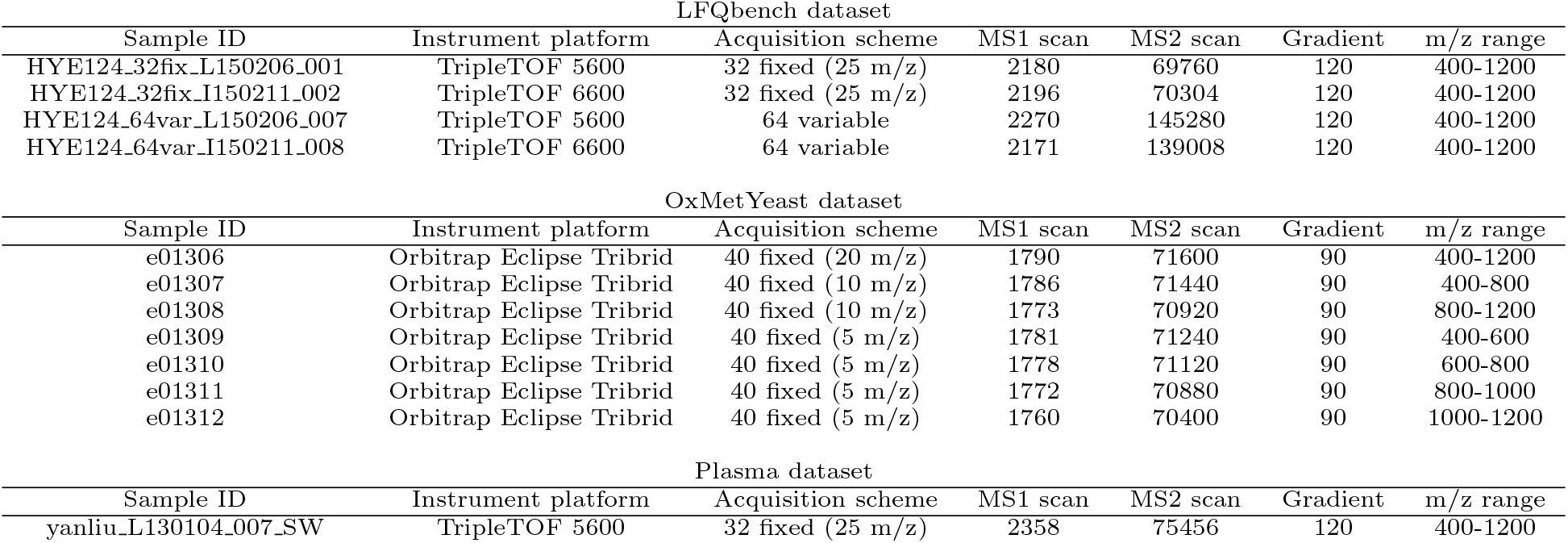
Properties of the three datasets—LFQbench, Plasma and OxMetYeast—used for evaluation.

**Table S2:** DIAmeter parameters

| DIAMeter |  |  |  |  |
| --- | --- | --- | --- | --- |
|  | Parameters | Orbitrap | TripleTOF 5600 | TripleTOF 6600 |
| Signal Extraction | k | 5 |  |  |
| tide-index | missed-cleavages | 2 |  |  |
|  | mods-spec | C+57.02146,3M+15.994915 |  |  |
|  | enzyme | trypsin |  |  |
| tide-search | precursor-window | 10 | 30 |  |
|  | precursor-window-type | ppm |  |  |
|  | mz-bin-width | 0.02 |  |  |
|  | mz-bin-offset | 0 |  |  |

**Table S3:**
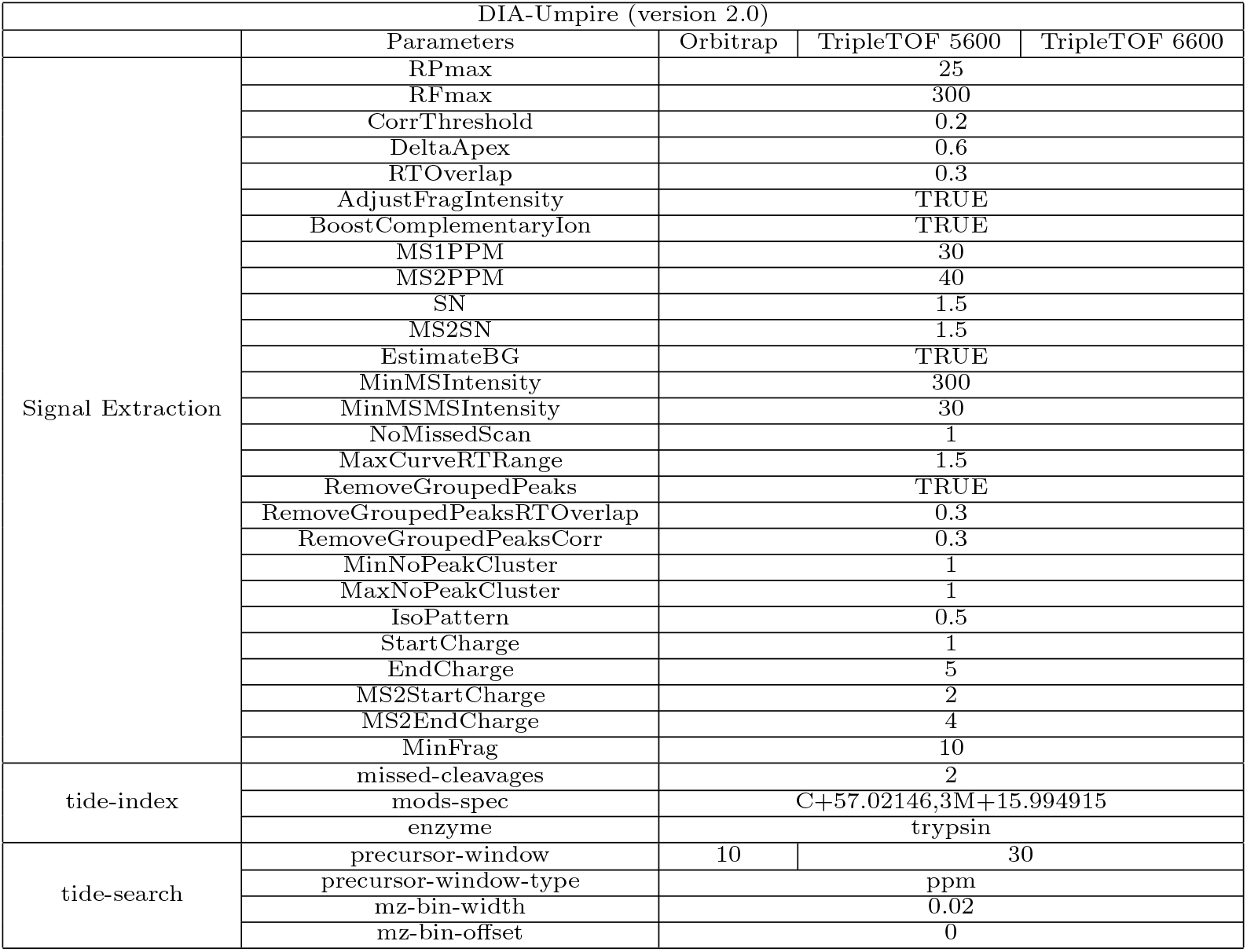
DIA-Umpire parameters.

**Table S4:** Parameters for Walnut (a re-implementation of PECAN in EncyclopeDIA^13^).

| PECAN / Walnut (version 0.9.0) |  |  |  |
| --- | --- | --- | --- |
| Parameters | Orbitrap | TripleTOF 5600 | TripleTOF 6600 |
| acquisition |  | DIA |  |
| filterPeaklists |  | TRUE |  |
| ptol | 10 | 30 |  |
| ftol | 10 | 30 |  |
| maxMissedCleavage |  | 2 |  |
| minCharge |  | 1 |  |
| maxCharge |  | 5 |  |

**Table S5:**
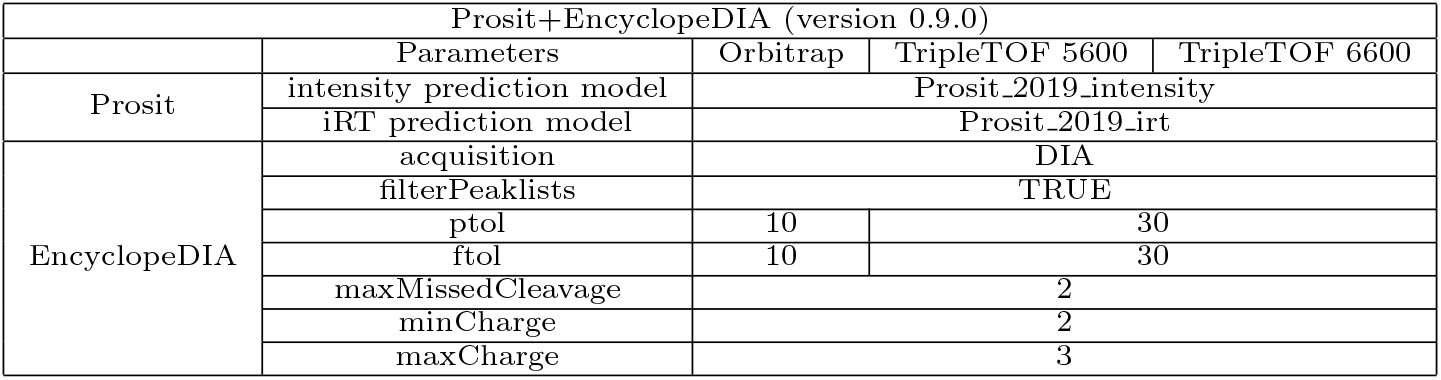
Prosit+EncyclopeDIA parameters.

## Footnotes

1 Submitted 23 Jan 2021, awaiting PRIDE ID.

